# Development of a ferritin-based nanoparticle vaccine against the SARS-CoV-2 Omicron variant

**DOI:** 10.1101/2022.03.13.484123

**Authors:** Wanbo Tai, Benjie Chai, Shengyong Feng, Xinyu Zhuang, Jun Ma, Mujia Pang, Lin Pan, Zi Yang, Mingyao Tian, Gong Cheng

**Author notes:** These authors contributed equally to this work (Wanbo Tai, Benjie Chai, Shengyong Feng). Corresponding to (G.C.).

## Abstract

A new SARS-CoV-2 variant named Omicron (B.1.1.529) discovered initially in South Africa has recently been proposed as a variant of concern (VOC) by the World Health Organization, because of its high transmissibility and resistance to current vaccines and therapeutic antibodies. Therefore, rapid development of vaccines against prevalent variants including Omicron is urgently needed for COVID-19 prevention. Here, we designed a self-assembling ferritin-based nanoparticle (FNP) vaccine against the SARS-CoV-2 Omicron variant. The purified Fc-RBD_Omicron_ automatically formed a dimer depending on the nature of the Fc tag, thus assembling onto the nanoparticles by the Fc-protein A tag interaction (FNP-Fc-RBD_Omicron_). The results of hACE2-transgenic mice immunization showed that SARS-CoV-2 Omicron RBD-specific IgG titer induced by FNP-Fc-RBDOmicron was much higher than that by Fc-RBD_Omicron_. Consistently, the sera showed a higher neutralizing activity against SARS-CoV-2 Omicron BA.1 and BA.2 in the FNP-Fc-RBD_Omicron_ immunized mice, indicating that immunization of a self-assembling ferritin-based nanoparticle vaccine offers a robust humoral immune response against Omicron variants. This study offers a great potential for the quick response of the emerging SARS-CoV-2 variants and affords versatility to develop universal vaccines against other emerging and reemerging coronaviruses in the future.

## Main Text

The COVID-19 pandemic has had a devastating effect on global health, resulting in over 6.0 million deaths worldwide. Continuous emergence of adaptive mutations of SARS-CoV-2 alters its pathogenicity and transmissibility, and renders its resistance to current vaccines and antiviral drugs.^1^ A new variant named Omicron (B.1.1.529) discovered initially in South Africa has recently been proposed as a variant of concern (VOC) by the World Health Organization, because of its high transmissibility and resistance to current vaccines and therapeutic antibodies.^2^ Therefore, rapid development of vaccines against prevalent variants including Omicron is urgently needed for COVID-19 prevention.

A previous study developed a SARS-CoV-2 vaccine based on a virus-like nanoparticle (VLP) platform, in which sixty copies of a fusion protein including a receptor binding domain (RBD) with a lumazine synthase as the structural scaffold were self-assembled into a nanoparticle.^3^ Immunization with this VLP-RBD conferred nearly complete protection to human ACE2 (hACE2)-transgenic mice against a high-dose SARS-CoV-2 infection.^4^ Based on this framework, we further designed a self-assembling ferritin-based nanoparticle (FNP) vaccine against the SARS-CoV-2 Omicron variant. In this system, twenty-four copies of ferritin containing an N-terminal protein A tag form a structural scaffold (Fig. 1a).^5^ The RBD (residues 331aa-524aa) of the SARS-CoV-2 Omicron spike protein with an Fc tag in the C-terminus (Fc-RBD_Omicron_) served as an essential immunogen (Fig. 1a).^6,7^ The purified Fc-RBD_Omicron_ automatically formed a dimer depending on the nature of the Fc tag^8^, thus assembling onto the nanoparticles by the Fc-protein A tag interaction (Fig. 1a). Based on this concept, the antigen of emerging SARS-CoV-2 variants can be rapidly assembled onto nanoparticles through a separating preparation and a subsequent Fc-Protein-A-tag-mediated conjugation.

**Fig. 1.**
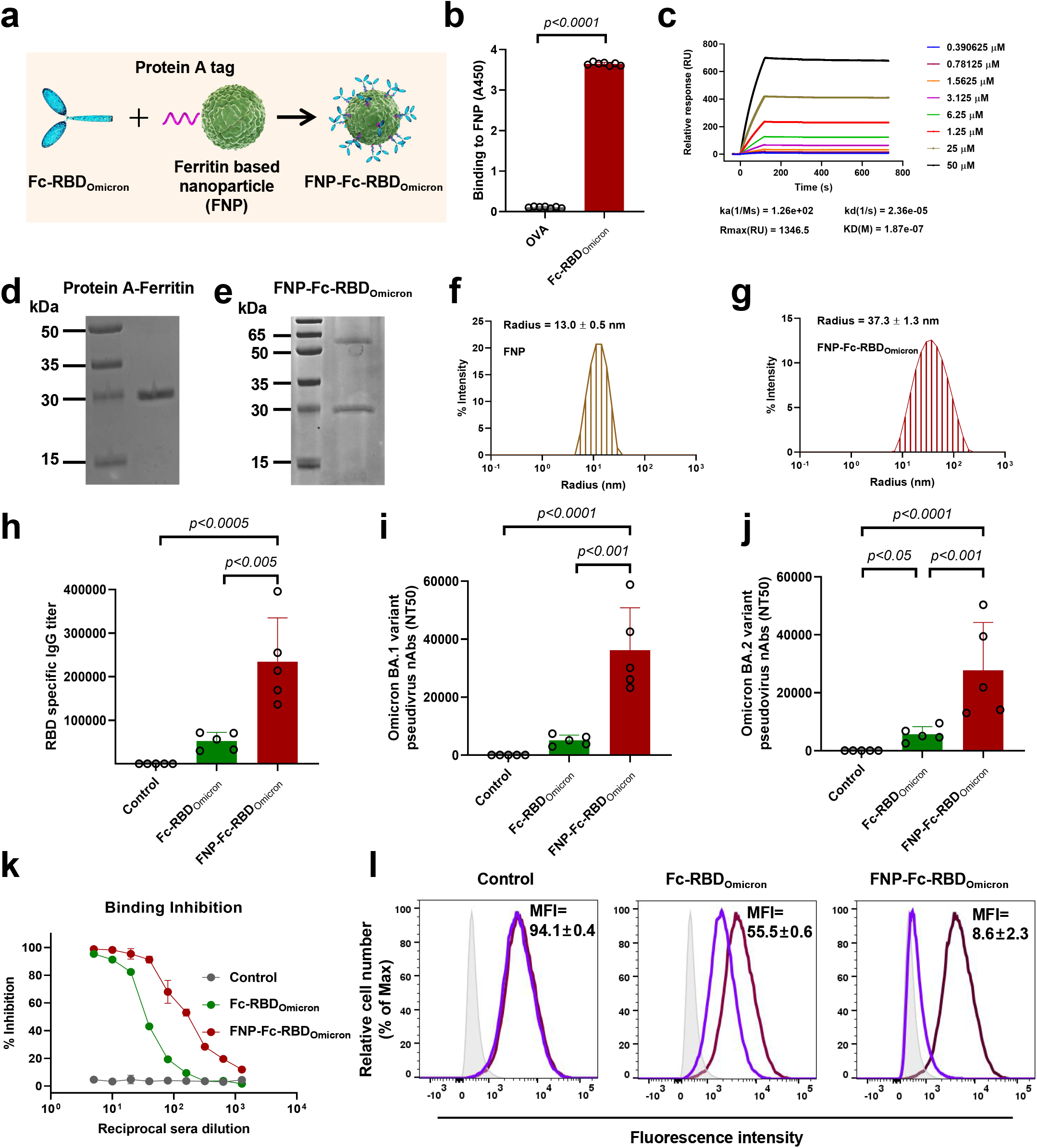
Development and characterization of the FNP-Fc-RBD_Omicron_ vaccine against SARS-CoV-2 Omicron variant. (a) Schematic representation of a SARS-CoV-2 Omicron RBD with Fc tag (light green), a ferritin-based 24-meric nanoparticle with N-terminal protein A tag (green), and an FNP-Fc-RBD_Omicron_ complex. (b-c) Interaction between Fc-RBD_Omicron_ and FNP was detected by ELISA (b) and by SPR (c). An equal amount of ovalbumin served as a control. Binding affinity was detected by an HRP-labeled anti-His tag antibody. The data are presented as mean±S.E.M. (n=7). (b) Statistical significance was calculated via ordinary unpaired parametric *t*-test. (d-e) The FNP complex with or without a Fc-RBD_Omicron_ was analyzed by SDS-PAGE. (f-g) Size distribution of the FNP complex with or without Fc-RBD_Omicorn_ was detected by DLS. (h-j) Measurement of IgG and neutralizing antibodies induced in immunized mice. The mice were immunized via intramuscular (i.m.) prime and boost at 2 weeks (10 μg per mouse, n=5). Sera at 14 days post-2nd immunization were detected for RBD_Omicron_-specific IgG antibodies by ELISA (h). The neutralizing antibodies were assessed by a pseudotyped SARS-CoV-2 Omicron BA.1 (i) and BA.2 (j) variants. The data are presented as mean±S.E.M. (n=5). Statistical significance was calculated via one-way ANOVA with multiple comparisons test. (k-l) Inhibition potency of immunized sera on SARS-CoV-2 RBD-hACE2 binding in hACE2/HEK293T cells. (k) The inhibition potency was evaluated by flow cytometry. Inhibition percentage (%) was calculated by a relative fluorescence intensity. (l) Representative images of the binding inhibition by the sera (1:40) from mice immunized with PBS (left panel), Fc-RBD_Omicron_ (middle panel), or FNP-Fc-RBD_Omicron_ (right panel). The violet lines represented median fluorescence intensity (MFI) values. The binding between Fc-RBD_Omicron_ and hACE2 is shown in dark red lines. Light gray shades indicate the Fc-hACE2 binding.

We expressed and purified the ferritin containing an N-terminal protein A tag in *Escherichia coli* (Fig. S1a), and its purity was confirmed by SDS-PAGE (Fig. 1d). The characterization of the self-assembling nanoparticles was analyzed by negative-stain electron microscopy (EM) (Fig. S1b) and dynamic light scattering (DLS) (Fig. 1f). The results indicated that the nanoparticles were spherical with a uniform diameter of 13.0±0.5 nm. We next expressed the Fc-RBD_Omicron_ in the FreeStyle 293-F cells (Fig. S2b). The binding affinity of Fc-RBD_Omicron_ for hACE2 was evaluated by both enzyme linked immunosorbent assay (ELISA) (Fig. S3b) and flow cytometry (Fig. S3c) with a dose-dependent manner. Incubation of Fc-RBD_Omicron_ with human HEK293T cells expressing human ACE2 (hACE2/HEK293T) potently interrupted the cellular entry of SARS-CoV-2 Omicron pseudoviruses, which were generated by a SARS-CoV-2 spike-pseudotyped human immunodeficiency virus (HIV) system.^9^ We next assembled the Fc-RBD_Omicron_ onto the 24-meric FNP by mixing these two components at a 24:1 molar ratio. The Fc-RBD_Omicron_ was capable of tightly interacting with the nanoparticles through the Fc-Protein A, measured by ELISA (Fig.1b) and surface plasmon resonance technology (SPR) (Fig.1c). The protein complex was co-eluted and co-purified by gel filtration chromatography, and further evaluated by SDS-PAGE (Fig.1e). The protein complex was designated as the FNP-Fc-RBD_Omicron_ throughout this investigation. Furthermore, the FNP-Fc-RBD_Omicron_ complex was evaluated by DLS, which confirmed the diameter of FNP-Fc-RBD_Omicron_ being uniformly about 37.3±1.3 nm (Fig.1g). Altogether, we generated self-assembling ferritin-based nanoparticles to develop a vaccine against the SARS-CoV-2 Omicron variant.

We next evaluated the potency of the FNP-Fc-RBD_Omicron_ to induce neutralizing antibody responses against SARS-CoV-2. To this end, we immunized hACE2-transgenic mice with either FNP-Fc-RBD_Omicron_ or a sole Fc-RBD_Omicron_. The mice were further boosted with the same dose of immunogens at 2 weeks after the primary immunization. Mouse sera were collected on Day 14 after the second immunization and analyzed for antibody titers and potency to neutralize SARS-CoV-2. The SARS-CoV-2 Omicron RBD-specific IgG titer induced by FNP-Fc-RBD_Omicron_ was 4 times higher than that by Fc-RBD_Omicron_ (Fig. 1h). Subsequently, the neutralizing potency in the sera of immunized animals was assessed by HIV pseudoviruses with SARS-CoV-2 Omicron BA.1 and BA.2 spikes. The sera showed a higher neutralizing activity in the FNP-Fc-RBD_Omicron_ immunized mice than that of a sole Fc-RBD_Omicron_ immunization (Fig.1i, 1j). To substantiate the SARS-CoV-2-neutralizing mechanism of vaccine-induced antibodies, we examined the interactions between the SARS-CoV-2 RBD and hACE2 in the presence of the vaccinated mouse sera by flow cytometry. Although the binding of RBD_Omicron_ to hACE2/HKE293T cells was inhibited by either FNP-Fc-RBD_Omicron_ or Fc-RBD_Omicron_ serum effectively in a dose-dependent manner, the former was more potent (Fig. 1k, 1l). We next examined whether the antibodies induced by FNP-Fc-RBD_Omicron_ immunization could interrupt the entry of HIV pseudotyped with other spike of SARS-CoV-2 VOCs. Indeed, FNP-Fc-RBD_Omicron_ vaccinated sera effectively blocked the cellular entry of multiple SARS-CoV-2 variants (Fig. S4), indicating that the Omicron RBD-based vaccine may be an ideal booster against all SARS-CoV-2 variants. Based on previous reports, neutralizing antibody titers induced by vaccines were well correlated with their *in vivo* protective efficacy against SARS-CoV-2 infection,^10^ thus the FNP-Fc-RBD_Omicron_ vaccine may be worth of preclinical testing in animals in the near future.

Overall, these results demonstrate that immunization of a self-assembling ferritin-based nanoparticle vaccine offers a robust humoral immune response against Omicron variant. Herein, a pseudovirus-based neutralization assay confirmed that vaccination with FNP-Fc-RBD_Omicron_ could provide an effective neutralizing potency against both Omicron BA.1 and BA.2 variant infection. This study offers a great potential for the quick response of the emerging SARS-CoV-2 variants and affords versatility to develop universal vaccines against other emerging and reemerging coronaviruses in the future.

## Acknowledgements

This work was supported by the Emergency Key Program of Guangzhou Laboratory (EKPG21-33) to G.C., the National Natural Science Foundation of China (32188101, 81961160737 and 31825001), the National Key Research and Development Plan of China (2021YFC2300200, 2020YFC1200104 and 2017ZX10304402).

## Author contributions

W.T., B.C., S.F., X.Z., J.M., M.P, L.P., Z.Y. and M.T. conducted the study and analyzed the data. G.C. designed and supervised the study, wrote and revised the manuscript.

