## Supplementary material for "Development of a ferritin-based nanoparticle vaccine against the SARS-CoV-2 Omicron variant"

**Supplementary information**

**Materials and Methods**

**Ethics statement**

All mouse related work was performed strictly in accordance with the guidance and recommendations in the Guide for the Care and Use of Laboratory Animals (National Research Council Institute for Laboratory Animal Research). Experiments were conducted under animal use protocols approved by the Institutional Animal Care and Use Committees at the Shenzhen Bay Laboratory and Tsinghua University.

**Cell lines and plasmids**

HEK293T cells (human embryonic kidney cells) was obtained from the American Type Culture Collection (ATCC) and cultured in Dulbecco's modified Eagle medium (DMEM, supplemented with 10% fetal bovine serum and 100 units/mL penicillin-streptomycin). hACE2/HEK293T cells were kindly given by professor Qiang Ding from Tsinghua University and cultured with the same condition as HEK293T cells. The FreeStyle 293-F Cells were purchased from Gibco and cultured in the FreeStyle 293 Expression Medium (Gibco) (Supplemented with 100 units/mL penicillin and 100 μg/mL streptomycin). BL21 (DE3) *E. coli* cells were obtained from TransGen Biotech and cultured in Luria-Bertani (LB) medium containing 50 μg/mL Kanamycin.

The genes encoding SARS-CoV-2 Omicron variant spike (GISAID accession number: EPI_ISL_2423556), ferritin (NCBI Reference Sequence: WP_000949190.1), and domain B of protein A (residues 212aa–270aa) from *S. aureus* (NCBI Reference Sequence: WP_190282922.1) were all optimized and synthesized by GenScript. The SARS-CoV-2 Omicron variant RBD (residues 331aa-524aa) was subcloned into pFuse-hIgG1-Fc2 vector (InvivoGen). Ferritin was subcloned into pET-28a c (+) vector with an N-terminal domain B of protein A and His×8-tag. Plasmids expressing the spike proteins of SARS-CoV-2 wild type and various variants pcDNA3.1-WT-S (Wuhan-Hu-1 strain, GenBank accession number: NC_045512.2) pcDNA3.1-B.1.1.7-S (Alpha variant, GISAID accession ID: EPI_ISL_601443), pcDNA3.1-B.1.351-S (Beta variant, GISAID accession ID: EPI_ISL_700428), pcDNA3.1-P.1-S (Gamma variant, GISAID accession ID: EPI_ISL_792680), pcDNA3.1-B.1.617-S (Delta variant, GISAID accession ID: EPI_ISL_2461258), pcDNA3.1-B.1.1.529-BA.1-S (Omicron variant, GISAID accession ID: EPI_ISL_6640916) and pcDNA3.1-B.1.1.529-BA.2-S (Omicron variant,

GenBank accession number: UJL09565.1) pNL4-3.luc.RE (the luciferase reporter-expressing HIV-1 backbone) were constructed previously and maintained in our laboratory.

**Protein expression and purification**

SARS-CoV-2 RBD protein was expressed from the FreeStyle 293-F cells. Briefly, the Fc-tagged RBD (Fc-$RBD_{Omicron}$) was collected from the cell culture medium, purified using Protein A column and Superdex 200 Increase 10/300 GL gel filtration chromatography. His-tagged ferritin-based nanoparticle (FNP) was expressed from BL21 (DE3) *E. coli* cells. Protein expression was induced using IPTG (isopropyl-beta-D-thiogalactoside) at a final concentration of 1 mM and purified using Ni-NTA column and Superdex 200 Increase 10/300 GL gel filtration chromatography. The purified FNP and Fc-$RBD_{Omicron}$ proteins were co-incubated (molar ratio is 1:24) at room temperature for 1 hour and subsequently the formed complex was purified using gel filtration chromatography. The diameters of FNP and FNP-Fc-$RBD_{Omicron}$ were characterized with a dynamic light scatter (DLS, Wyatt Technology), and the purified proteins were analyzed by SDS-PAGE.

**Surface plasmon resonance (SPR) analysis**

The binding kinetics of Fc-$RBD_{Omicron}$ with FNP were analyzed by SPR (Biacore 8K, GE Healthcare). Specifically, the FNP were transferred into HBST buffer (20 mM HEPES (pH 7.4), 150 mM NaCl, and 0.005% (v/v) Tween 20) and immobilized on the CM5 chip (Cytiva). Then, serially diluted Fc-$RBD_{Omicron}$ samples (from 50 μM to 0.390625 μM) flowed over the chip in PBST buffer. BSA protein was selected as negative control. Binding affinities were measured using a BIAcore 8K (GE Healthcare) at 25°C in the multi-cycle mode. Binding kinetics were analyzed with Biacore$^{TM}$ Insight software (GE healthcare) using a 1:1 Langmuir binding model.

**Negative staining analysis**

Negative-staining electron microscopy procedures were conducted as previously described.[1] Briefly, the purified FNP sample of 5 μl with a final concentration of 0.15 mg/ml in PBS was loaded onto a freshly glow-discharged carbon coated grid (230 mesh, Beijing Zhongjingkeyi). After incubating for 1 min, excess sample was blotted, and the grid was stained with 5 μl 2% (w/v) uranyl acetate solution for 1 min. Excess solution was blotted and grids were dried at room temperature. Images were acquired using a Tecnai Spirit (FEI) operated at 120 kV and 4 K × 4 K Ultrascan CCD camera at 98,000× magnification at the Institute of Biophysics, Chinese Academy of Sciences.

**Mouse immunization**

Four-week-old hACE2 transgenic mice were immunized with FNP-Fc-$RBD_{Omicron}$ protein (10 μg/mouse), Fc-$RBD_{Omicron}$ protein (10 μg/mouse), or PBS control in the presence of aluminum adjuvants (500 μg/mouse, InvivoGen) via intramuscular route. The immunized mice were boosted 14 days later with the same dose immunogen and adjuvants, and sera were collected at 14 days after the 2nd immunizations for specific IgG antibodies or neutralizing antibodies analysis.

**ELISA**

ELISA was carried out to detect the binding of Fc-$RBD_{Omicron}$ to soluble hACE2 protein, and human IgG Fc (Fc, Thermo Fisher Scientific) protein were used as control. Briefly, ELISA plates were precoated with SARS-CoV-2 RBD or Fc protein (1 μg/ml) overnight at 4 °C and blocked with 2% fat-free milk in PBST for 2 h at 37 °C. Serially diluted $His_{\times 6}$ tagged hACE2 protein (Sino Biological) was added to the plates and incubated for 2 h at 37 °C. After four washes, the bound protein was detected using anti-His tag antibody (HRP) (0.005 μg/ml, Sino Biological) for 1 h at room temperature. The reaction was visualized by addition of substrate 3,3',5,5'-Tetramethylbenzidine (TMB, Sigma) and stopped by $H_2SO_4$ (1 N). The absorbance at 450 nm was measured by an ELISA plate reader.

Next, ELISA was conducted to detect the binding of SARS-CoV-2 Fc-$RBD_{Omicron}$ protein to FNP, and ovalbumin (OVA, Invivogen) was set as controls. Like the above description, ELISA plates were precoated with Fc-$RBD_{Omicron}$ or OVA at 1 μg/ml. And then $His_{\times 8}$ tagged FNP protein (0.5 μg/ml) was added to the wells and incubated for 2 h at 37 °C. After four washes, the binding was detected using HRP labeled anti-His tag antibody (0.005 μg/ml, Sino Biological) for 1 h at room temperature. The reaction was visualized by addition of TMB (Sigma) and stopped by $H_2SO_4$ (1 N). The absorbance at 450 nm was measured by an ELISA plate reader.

ELISA was also performed to detect the interaction between SARS-CoV-2 Omicron RBD protein and RBD-specific antibodies in mouse sera. The procedure was the same as described above, except that the ELISA plates were coated with RBD (Sino Biological) at 1 μg/ml and then sequentially incubated with serially diluted mouse sera and HRP-conjugated anti-mouse antibodies (1:5000, Thermo Fisher Scientific).

**Pseudovirus neutralization and inhibition assays**

SARS-CoV-2 pseudovirus was generated, as previously described.[2] Briefly, HEK293T cells were cotransfected with a plasmid encoding Env-defective, luciferase-expressing HIV-1 genome (pNL4-3.luc.RE) and a plasmid encoding SARS-CoV-2 S protein using the calcium phosphate method. The transfected medium was replaced by fresh DMEM 8 h later, and pseudovirus-containing supernatants were collected 48 h later for single-cycle infection in hACE2/HEK293T cells. Pseudovirus neutralization assay was then performed by incubation of SARS-CoV-2 pseudovirus with serially diluted mouse sera for 1 h at 37 °C, followed by addition of the mixture into hACE2/HEK293T cells. Fresh medium was added 24 h later, and the cells were lysed 72 h later in cell lysis buffer (Promega). The lysed cell supernatants were incubated with luciferase substrate (Promega) and detected for relative luciferase activity. The 50% pseudovirus neutralizing antibody titer ($NT_{50}$) was calculated.

Pseudovirus entry inhibition by Fc-$RBD_{Omicron}$ was next carried out. Briefly, Fc-$RBD_{Omicron}$ protein at serial dilutions was incubated with hACE2/HEK293T target cells for 1 h at 37 °C. The cells were then infected with SARS-CoV-2 pseudovirus Omicron variant. The Fc protein was used as controls. Fresh medium was added 24 h later, and the cells were lysed and analyzed as described above.

**Flow cytometry**

Flow cytometry analysis was first performed to detect the binding of the Fc-$RBD_{Omicron}$ protein to hACE2/HEK293T cell. The Fc protein was used as controls. Briefly, cells were incubated with reciprocal diluted Fc-$RBD_{Omicron}$ for 30 min at room temperature, after three washes with PBS (containing 2% FBS), the cells were incubated with FITC-labeled goat anti-human IgG antibody (1:500, Thermo Fisher Scientific) at room temperature for 20 min. After washes, the cells were fixed with 4% formaldehyde and the fluorescence intensity of the cells was measured using flow cytometry (BD LSRFortessa$^{TM}$ system).

Flow cytometry analysis was next conducted to detect the interaction between the Fc-$RBD_{Omicron}$ and hACE2 in the presence of mouse sera. Briefly, hACE2/HEK293T cells were incubated with Fc-$RBD_{Omicron}$ (10 μg/ml) in the presence or absence of serially diluted mouse sera at room temperature for 1 h, which was followed by incubation with FITC-labeled goat anti-human IgG antibody (1:500, Thermo Fisher Scientific) for 30 min and analyzed.

**Statistical analysis**

The results are presented as the mean±standard error of mean (S.E.M.). The difference between any two groups were determined by unpaired parametric *t*-test or one-way ANOVA with multiple comparisons tests depending on the distribution of the data. All data were analyzed with GraphPad Prism version 8.0 software.

**Supplementary Figures**

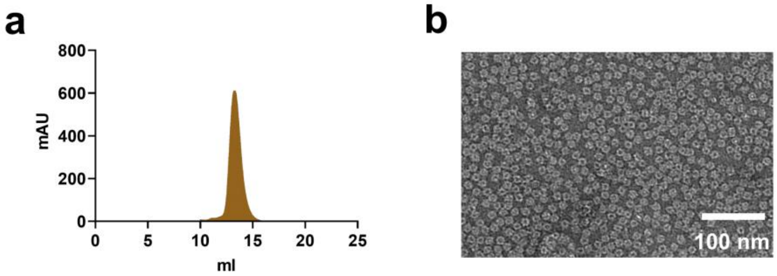


**Fig. S1 Preparation and characterization of 24-meric ferritin-based nanoparticle.**

The ferritin with an N-terminal protein A tag was expressed in BL21 (DE3) *E. coli* cells, and then purified to high quality. (a) Representative elution profiles of the nanoparticles from Superdex 200 Increase 10/300 GL gel filtration chromatography recorded at milli-absorbance unit (mAu) at 280 nm wavelength. (b) Negative-staining EM analysis of the nanoparticles. Scale bar, 100 µm.

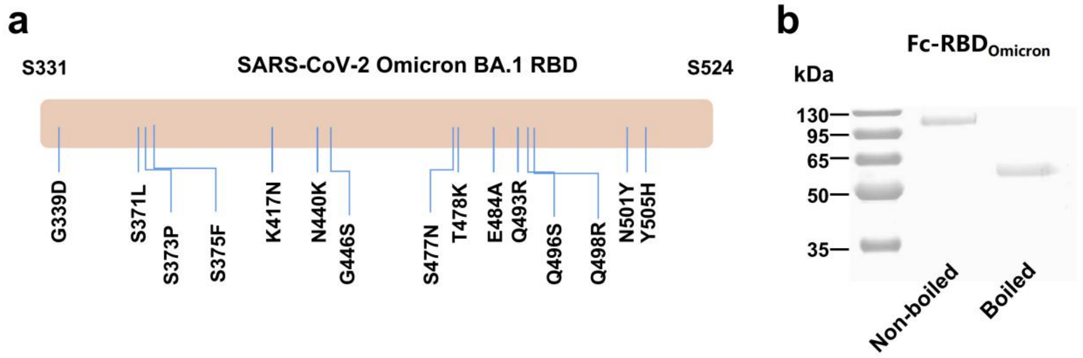


**Fig. S2 Characterization of the Fc tagged RBD of SARS-CoV-2 Omicron variant.**

(a) The schematic diagram showing the RBD (residues 331aa-524aa of the spike) mutations of the SARS-CoV-2 Omicron variant. Compared to the Wuhan-Hu-1 strain, total 15 mutations were included and labeled. (b) Expression of Fc-$RBD_{Omicron}$ protein in the FreeStyle 293-F cells. Cells were transfected with Omicron RBD-encoding plasmid, and the supernatant was purified with protein A beads and gel filtration chromatography at 72 h after transfection. and then the boiled or non-boiled Fc-$RBD_{Omicorn}$ samples was analyzed by SDS-PAGE under reducing condition, the molecular weight markers were indicated on the left.

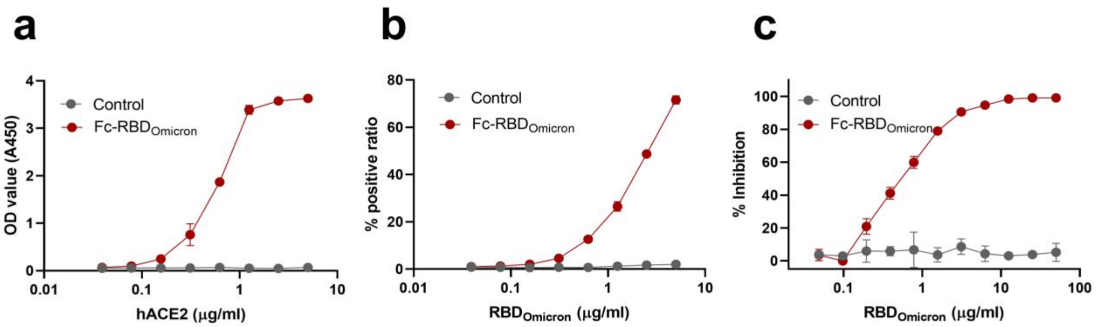


**Fig. S3 Incubation of the Fc-RBD$_{Omicron}$ interrupted the hACE2-mediated SARS-CoV-2 entry.**

(a) Detection of Fc-RBD$_{Omicron}$ binding to hACE2 by ELISA. The data are presented as mean±S.E.M. (n=3). (b) Binding of Fc-RBD$_{Omicron}$ protein to hACE2/HEK293T cells by flow cytometry. The data are presented as mean±S.E.M. (n=3). (c) Inhibition of Fc-RBD$_{Omicron}$ against pseudotyped SARS-CoV-2 Omicron variant entry into hACE2/HEK293T cells. The data are presented as mean±S.E.M. (n=3). Human IgG Fc protein acted as a control. Experiments were repeated twice with similar results.

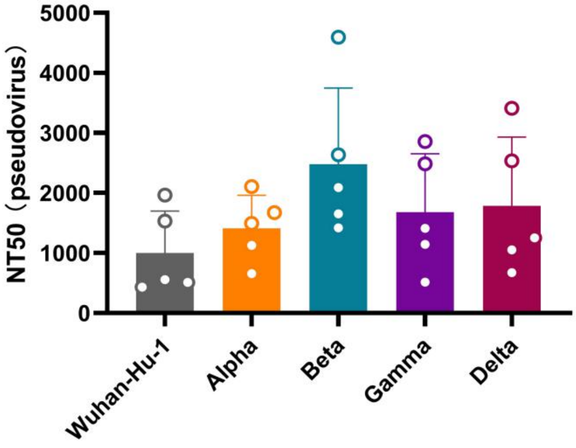


**Fig. S4 Antibodies induced by the FNP-Fc-RBD$_{\mathbf{Omicron}}$ immunization neutralize pseudotyped SARS-CoV-2 variants.**

The cross-neutralizing antibodies from FNP-Fc-RBD$_{Omicron}$ immunized sera (3-fold serial dilutions from 1:50) were assessed to interrupt the cellular entry of pseudoviruses of SARS-CoV-2 VOCs (Wuhan-Hu-1, Alpha, Beta, Gamma and Delta) in hACE2/HEK293T cells. The data are presented as mean±S.E.M. (n=5). Experiments were repeated twice with similar results.
